# Prenatal heat stress effects on gestation and postnatal behavior in kid goats

**DOI:** 10.1101/701151

**Authors:** Wellington Coloma-García, Nabil Mehaba, Pol Llonch, Gerardo Caja, Xavier Such, Ahmed A. K. Salama

## Abstract

Consequences of heat stress during pregnancy can affect the normal development of the offspring. In the present experiment, 30 Murciano-Granadina dairy goats (41.8 ± 5.7 kg) were exposed to 2 thermal environments varying in temperature-humidity index (THI) from 12 days before mating to 45 days of gestation. The environmental conditions were: gestation thermal-neutral (GTN; THI = 71 ± 3); and gestation heat stress (GHS; THI = 85 ± 3). At 27 ± 4 days old, GTN-born female kids (n = 16) and GHS-born ones (n = 10) were subjected to 2 tests: arena test (AT) and novel object test (NOT), the latter was repeated at 3 months of age. Additionally, 8 months after birth, a subset of growing goats (n = 8) coming from GTN and GHS (16.8 ± 3.4 kg BW) were exposed consecutively to 2 environmental conditions: a basal thermal-neutral period (THI = 72 ± 3) for 7 days, and a heat-stress period (THI = 87 ± 2) for 21 days. In both periods, feeding behavior, resting behavior, other active behaviors (exploring, grooming), thermally-associated behaviors and posture were recorded. The gestation length was shortened by 3 days in GHS goats. In the AT, GHS kids showed a lower number of sniffs (*P* < 0.01) compared to GTN. In the NOT, GHS kids also tended to show a lower number of sniffs (*P* = 0.09). During heat exposure, GTN and GHS growing goats spent more time resting as well as exhibited more heat-stress related behaviors such as panting and drinking (*P* < 0.001); however, no differences were observed between both groups. In conclusion, heat stress during the first third of pregnancy shortened gestation length and influenced the exploratory behavior of the kids in the early life without impact on the behavior during the adulthood when exposed to heat stress.

## Introduction

There is evidence that environmental conditions during pregnancy can modify fetal programming through physiological and epigenetic changes [1, 2], which permanently modify the behavior, health and productivity of the offspring. Several studies have shown that episodes of stress during the prenatal stage have negative effects on the pregnancy itself, by shortening its duration [3, 4], and on the postnatal life of the offspring by reducing birth weight [2].

Beyond these effects, maternal stress during pregnancy has shown to have profound effects on the development and function of the hypothalamic-pituitary-adrenal (HPA) axis, and the associated circulating ACTH and cortisol concentrations [5]. Moreover, recent research suggests that these effects remain in further generations [6]. In this regard, most studies using rodent or primate models, have evidenced the negative impact of gestational stress on increased aggressiveness and altered social interactions [7-9] and a reduction in the neuromotor capacities and exploration and learning [4,10].

In the future, as global warming progresses, an increase in temperatures accompanied by increasingly frequent heat waves is expected [11]. In the case of ruminants, heat stress during pregnancy has attracted special attention, due to the significant impact on food production (i.e. milk) [12]. Furthermore, although literature is scarce, thermal stress during pregnancy is demonstrated to be responsible for the abnormal development of the fetus and cause a harmful effect in the early postpartum period and adulthood. For instance, prenatal heat stress can impair the normal postnatal growth of the offspring and compromise their passive immunity but also alter the behavioral patterns [13,14]. Nevertheless, the previous studies evaluated the impact of maternal heat stress during the late gestation in cows, but little is known about the effects of heat stress during early pregnancy on offspring behavior in dairy animals, including cows and goats. There is strong evidence that fetal programming occurs during early gestation in ruminants, and several environmental and nutritional factors during this period can condition performance of offspring permanently. For instance, adequate maternal nutrition in early gestation is critical for the normal development of all fetal organs and tissues [15]. Additionally, exposure of cows to limited nutrition during early gestation resulted in decreased skeletal muscle mass and altered glucose metabolism of offspring [16]. Therefore, we hypothesized that heat stress (with its related effects such as altered blood flow, changes in hormone levels, reduced feed intake, etc.) during early gestation would alter performance and response of offspring to environmental stimuli.

Behavior is a phenotypical trait that is very sensitive to the environment. One of the first changes that can be observed in animals that are under stressful conditions is a change in their behavior repertoire. Within behavior, the way animals react to novel situations is also influenced by the environmental conditions in where they live [17]. Therefore, behavior is a sensitive measure to investigate changes of perception of the environment. In the present study, it was investigated the effect of heat stress in goats at the beginning of the pregnancy on the gestation performance and the changes in the behavior of the offspring when challenged with heat, both at neonatal and adult stages.

## Materials and methods

The animal care conditions, treatment, housing, and management practices followed the procedures stated by the Ethical Committee of Animals and Humans Experimentation of the UAB (4790) and following the EU legislation (Regulation 2010/63/EC).

### Treatments and management conditions of dams

Thirty multiparous lactating Murciano-Granadina dairy goats of 41.8 ± 5.7 kg body weight (BW) bred at the experimental farm of UAB were used. Goats were housed in 6 pens (5 × 2.5 m^2^) of 5 goats each, distributed equally in 2 adjacent rooms, one for each treatment. Goats were distributed by similar BW within each pen. The present experiment was carried out during spring (March to June). After 2 weeks of adaptation to the experimental conditions, goats were distributed in 2 groups exposed to 2 different climatic conditions (n = 15) from day 12 before mating until day 45 of gestation. The climatic conditions were: thermo-neutral (TN), and heat stress conditions (HS). The TN group was maintained between 15 and 20°C (room temperature), and 49 ± 8% relative humidity (temperature humidity index, THI = 71 ± 3, calculated according to NRC [18]), and HS group for 12-h day at 37 ± 0.5°C and 45 ± 5% relative humidity (THI = 85 ± 3) and 12-h night at 30 ± 0.5°C and 47 ± 2% relative humidity (THI = 80 ± 2). The room housing HS animals was equipped with 4 electric heaters coupled to thermostat (3.5 kW; General Electric, Barcelona, Spain). Environmental temperature and humidity were recorded every 10 min throughout the experiment by data loggers (Opus 10, Lufft, Fellbach, Germany). Both treatments were maintained from 12 days before mating until 45 days after mating (early gestation).

Estrus was synchronized in 2-day intervals. Synchronization was performed using intravaginal sponges (progesterone P4; Sincropart 30 mg, Ceva Animal Health, Barcelona, Spain) for 12 days followed by the administration of equine-chorionic gonadotropin (eCG, 400 IU; Ceva Animal Health) at the time of sponge withdrawal. Goats were naturally mated by the same buck at 2-day intervals.

Feed was provided *ad libitum* as a total mixed ration (70% alfalfa hay and 30% concentrate). Concentrate contained barley 31.5%, corn 41.5%, soybean meal 44.5%, sodium bicarbonate 1%, calcium phosphate 0.4%, calcium carbonate 0.5%, salt 0.7%, and premix 0.4%; as fed basis. Water was freely available at room temperature. Mineral salt blocks (Na 36.7%, Ca 0.32%, Mg 1.09%, Zn 5 g/kg, Mn 1.5g /kg, S 912 mg/kg, Fe 304 mg/kg, I 75 mg/kg, Co 50 mg/kg, and Se 25 mg/kg; Ovi Bloc, Sal Cupido, Terrassa, Spain) were freely available in each pen throughout the experiment.

Goats were milked twice per day using a mobile milking unit set at 42 kPa, 90 pulses/min, and 66% pulsation ratio. Feed intake was recorded daily, calculated by the difference between the weight of the ration offered and the leftover at the end of the day. Rectal temperature (RT) and respiration rate (RR) were recorded daily 3 times per day at 8, 12, and 17 h. RT was recorded with a digital veterinary thermometer (ST714AC Accu-vet, Tecnovet S.L, Barcelona, Spain). RR was calculated as the number of breaths per minute by counting the flank movements with the help of a chronograph and from a distance of 2 m without disturbing the goats.

Pregnancy was confirmed by trans-rectal ultrasound at day 21 and 45 after mating, and all goats were confirmed to be pregnant. After 45 days of gestation, all goats were gathered in one group and managed under semi-intensive conditions (grazing 6 h/day and feed complemented when indoors). Two weeks before the expected date of parturition, the goats were weighed and moved to kidding pens for permanent surveillance and parturition assistance. Immediately after birth, kids were separated from the goats and fed with their mothers’ colostrum and reared together with milk replacer (150 g/L, Elvor, Saint-Brice, France) with an automatic milk provider (Foerster-technik, Engen, Germany). Pregnancy length and litter size of kids were recorded after parturition. BW of kids was recorded at birth and every week until 4 weeks old with a digital scale (Tru-Test AG500 Digital Indicator, accuracy, Auckland, New Zealand).

### Behavioral tests and measurements with kids

For the behavioral assessment, female kids at 27 ± 4 days old, from gestation TN (GTN; n = (16) or gestation HS (GHS; n = 10) conditions were individually exposed to an arena test (AT) for 5 consecutive days, and to a novel object test (NOT) at 48 h after the end of the AT. The NOT test was repeated at 3 months of age. Behavioral tests were carried out into an artificial climatic chamber (Eurosheild, ETS Lindgren-Euroshield Oy, Eura, Finland) in order to avoid sounds from outside and variations of temperature. All tests were video recorded for subsequent analysis.

### Arena test (AT)

The AT was carried out in a 4 × 4 × 2.3 m^3^ arena (w × l × h), in which 9 squares of 1.3 × 1.3 m^2^ were painted on the ground with chalk. The access to the arena was through a starting cage of 50 × 50 × 60 cm^3^ (w × l × h) separated from the arena by a guillotine door (Fig A in S1). On the test day, each kid was randomly selected among the 2 treatments, transported from the nursery to the starting cage and freed 30 s later into the arena. The duration of the test was 8 min and time started to run when the kid was completely inside the arena. The following behavioral parameters were measured: number of squares entered, frequency of jumping and sniffing (nose less than 5 cm from the walls or floor) events, number of vocalizations and distance walked (movement forward).

### Novel object test (NOT)

With NOT the same procedure was followed as for AT and the same behavioral measurements were registered. In addition, a road hazard cone (0.5 × 0.7 m^2^, w × h) was placed on the floor against the wall opposite to the starting cage (Fig B in S1), thereby the latency and the frequency of sniffing events addressed to the novel object were registered. The NOT test was repeated one month later.

### Heat-stress challenge trial with growing goats

To compare the behavioral response of animals born from GTN and GHS goats to the same stressor (i.e., heat stress) after sexual maturity, a subset of the growing goats was selected at 8 months of age. The animals were balanced by BW and mother parity, and randomly allocated to individual pens (1.08 m^2^) with 8 replicates per group. After one week for adaptation to facilities, 2 different climatic conditions were applied in 2 consecutive periods to both groups, following a randomized controlled design. During the first period, basal period (1 week), temperature and humidity were in average 24 ± 2.43°C and 68 ± 9% (THI = 72), respectively. On the other hand, during the heat-stress challenge period (3 weeks), the average temperature was 37 ± 1.8°C and humidity was 49 ± 7.0% (THI = 87) during the day and 31 ± 1.4°C and 53 ± 7.0% (THI = 80), respectively, at night. Room temperature was automatically controlled with a thermostat (3.5 kW; General Electric, Barcelona, Spain) regulating 4 electric heaters. Environmental temperature and humidity were continuously recorded every 10 min throughout the experiment by data loggers (Opus 10, Lufft, Fellbach, Germany).

Feed was provided as a total mixed ration consisting of 85% alfalfa hay and 15% concentrate (as feed basis: oat grain 5%, malting barley 10%, canola meal 10%, gluten feed 10%, corn 4.7 %, soy hulls 45%, soybean oil 5%, soybean meal 5%, molasses 2%, bicalcic phosphate 2.5%, salt 0.5%, premix 0.3%) once daily at 9:30 h. Clean water was freely and individually available for each goat.

### Behavior measurements of growing goats

A single trained observer recorded behavior following a scan-sampling methodology [19]. Behaviors were recorded between 12 h and 17 h, within the period of heat stress. The behavioral observations were performed daily and the duration of each session was 2 h, whereby each pen was scanned 40 times at 3 min interval.

The behavioral measurements were drawn from the Welfare Assessment Protocol for Goats [20]. Feeding (feeding + rumination + drinking), other non-feeding active and inactive behaviors (exploration + grooming + other + resting) and physiological behavior associated to thermal stress (open-mouth or close-mouth panting) were recorded as well as posture (standing-walking + standing-immobile + lying-straight + lying-joint). The definition of the recorded behaviors is presented in Table S1.

### Statistical analyses

The duration of pregnancy and birth weight were analyzed with the GLM procedure of SAS (version 9.4; SAS Institute Inc., Cary, NC). The feed intake and RT and RR measurements of the goats (dams) were analyzed as repeated measures using a linear mixed model (PROC MIXED procedure). Behavioral data from NAT and scan sampling during the heat exposure trial, as counts and week average percentages, respectively, were analyzed as repeated measures using a generalized linear mixed occasional behaviors, using a generalized linear model (PROC GENMOD), all adjusted under a Poisson or a Negative Binomial distribution, according to the fitness of the model. Also, litter size was analyzed using the PROC GENMOD procedure. The models included treatment (GHS vs GTN) as fixed effect, and in the case of repeated measures, day or week was also included as a fixed effect as well as the interaction of treatment × day or treatment × week, while animal was considered as a random effect. Litter size was also used as a covariable for the analysis of the duration of pregnancy and litter weight. Differences between least squares means were determined with the PDIFF test of SAS. Significance was declared at *P* < 0.05 and trend at *P* < 0.10 unless otherwise indicated.

## Results

### Effects of heat stress during the pregnancy and early postpartum

Regarding the physiology measurements of goats during the experimental period, GHS goats showed a higher RT compared to GTN goats (average 38.7°C for GTN and 39.3°C for GHS; SED = 0.07 and *P* < 0.01) and higher RR (average 33 breaths/min for GTN and 108 breaths/min for GHS; SED = 3.06 and *P* < 0.01), indicating that GHS treatment effectively triggered a heat stress response. Feed intake was lower in GHS compared to GTN goats (2.52 kg/day for GTN and 2.12 kg/day for GHS; SED = 0.55 and *P* = 0.001).

The results of the different variables evaluated at parturition and early postpartum period are shown in Table 1. The gestation length was on average shortened 3 days in GHS goats compared to GTN (*P* = 0.006). The litter weight of GHS group tended to be lower compared to GTN (*P* = 0.061). Litter weight showed to be influenced by the litter size (*P* < 0.001), as a greater litter size was associated to smaller kids. However, litter size and kids weight at 35 days of age were not affected by the treatment (*P* > 0.10).

**Table 1.** Gestation length in dams and performance of kids at birth and early postpartum period.

| Item | Treatment<sup>1</sup> | | SED<sup>2</sup> | Effect ($P$-value) | |
| --- | --- | --- | --- | --- | --- |
|  | GTN | GHS |  | Treatment | Litter size<sup>3</sup> |
| Litter size | 2.31 | 2.23 | 0.31 | 0.806 | - |
| Litter weight, kg | 5.40 | 4.71 | 0.71 | 0.061 | 0.001 |
| Duration of pregnancy, day | 146 | 143 | 0.9 | 0.006 | 0.915 |
| Birth-weight of kids<sup>4</sup>, kg | 2.34 | 2.18 | 0.10 | 0.122 | - |
| Weight of 35-days-old kids<sup>5</sup>, kg | 7.88 | 7.64 | 0.54 | 0.520 | - |

<sup>1</sup> GTN, dams exposed to thermal-neutral conditions during the first 45 days of gestation (n = 15); GHS, dams exposed to heat-stress during the first 45 days of gestation (n = 15).
<sup>2</sup> Standard error of the difference.
<sup>3</sup> Litter size used as a covariable.
<sup>4</sup> n = 30 kids for GTN, and n = 30 for GHS.
<sup>5</sup> n = 26 kids for GTN, and n = 23 for GHS.

### Behavioral tests on kids

The results of the behavioral tests are summarized in Table 2. In the arena test (AT), a significant day effect was observed in all parameters, as the number of vocalizations (*P* < 0.001) and walked distance (*P* = 0.001) decreased, whereas the number of jumping (*P* < 0.001) and sniffing events (*P* = 0.009) increased from day 1 to day 5, reflecting habituation of kids to the arena test facilities. Also, the number of squares that kids walked through showed to be higher from day 1 to day 2 afterwards being diminished towards day 5 (*P* ≤ 0.001), which is consistent with the reduction in the walked distance. Regarding the effect of treatment, GHS kids showed a lower number of sniffing events compared to GTN kids (*P* = 0.009). Additionally, the significant interaction between treatment and day for vocalizations (*P* < 0.001) was due to the fact that the number of vocalizations in the GHS kids was lower during the 2 first days (*P* ≤ 0.05) and recovered thereafter. The rest of behavioral parameters assessed were not influenced by the gestational exposure to heat stress (*P* > 0.10).

**Table 2.** Behavioral responses in arena test (AT) of female kids during 5 consecutive days.

| Item | Treatment<sup>1</sup> | | SED<sup>2</sup> | Effect ($P$-value) | | |
| --- | --- | --- | --- | --- | --- | --- |
|  | GTN | GHS |  | Trt<sup>3</sup> | Day | Trt×Day |
| No. of squares entered | 43.4 | 31.5 | 4.87 | 0.115 | 0.009 | 0.704 |
| No. of jumps | 1.54 | 1.15 | 0.476 | 0.586 | 0.001 | 0.546 |
| No. of sniffs of the arena | 33.5 | 26.7 | 1.62 | 0.007 | 0.001 | 0.335 |
| **No. of vocalizations** | 171 | 150 | 11.7 | 0.200 | 0.001 | 0.001 |
| **Distance travel (s)** | 54.8 | 44.9 | 7.06 | 0.282 | 0.001 | 0.123 |

<sup>1</sup> GTN, kids born to dams exposed to thermal-neutral conditions during the first 45 days of gestation (n = 16); GHS, kids born to dams exposed to heat stress conditions during the first 45 days of gestation (n = 10).
<sup>2</sup> Standard error of the difference.
<sup>3</sup> Trt, treatment effect (GHS vs GTN).

Regarding the novel object test (NOT) results (Table 3), this test was performed at 1 and 3 months of age. At 1 month of age, GHS kids showed a trend to reduce the number of sniffing events compared to GTN kids (*P* = 0.093) revealing a smaller motivation for exploration of novel objects in kids whose mothers suffered from heat stress during gestation. No treatment differences were detected in the rest of the behavioral parameters measured. At 3 months of age, no treatment effects were found on any of the parameters assessed in the NOT.

**Table 3.** Behavioral responses in novel arena test (NOT) of female kids at 1 and 3 months of age. Values are presented as means ± standard deviation.

| Item | Treatment<sup>1</sup> | | Effect ($P$-value) |
| --- | --- | --- | --- |
|  | GTN | GHS | Treatment |
| **1 month of age** |  |  |  |
| No. of squares entered | 47.5 ± 1.08 | 38.9 ± 1.10 | 0.127 |
| No. of jumps | 4.81 ± 1.700 | 2.30 ± 1.980 | 0.413 |
| No. of sniffs of the arena | 36.1 ± 1.06 | 30.3 ± 1.08 | 0.093 |
| No. of vocalizations | 156 ± 1.1 | 162 ± 1.1 | 0.670 |
| Distance travel (s) | 48.8 ± 3.87 | 41.0 ± 4.90 | 0.220 |
| No. of sniffs of the object | 14.8 ± 1.14 | 10.5 ± 1.19 | 0.136 |
| Latency before 1st sniff of the object (s) | 53.9 ± 53.80 | 77.4 ± 24.80 | 0.562 |
| **3 months of age** |  |  |  |
| No. of squares entered | 41.3 ± 1.10 | 41.3 ± 1.13 | 0.998 |
| No. of jumps | 0.31 ± 1.560 | 0.50 ± 1.560 | 0.461 |
| No. of sniffs of the arena | 30.1 ± 1.06 | 33.5 ± 1.08 | 0.286 |
| No. of vocalizations | 168 ± 1.0 | 157 ± 1.0 | 0.670 |
| Distance travel (s) | 59.0 ± 6.46 | 53.0 ± 8.17 | 0.609 |
| No. of sniffs of the object | 5.25 ± 1.150 | 4.40 ± 1.220 | 0.136 |
| Latency before 1st sniff of the object (s) | 43.2 ± 7.47 | 40.9 ± 9.96 | 0.855 |

<sup>1</sup> GTN, kids born to dams exposed to thermal-neutral conditions during the first 45 days of gestation (n = 16); GHS, kids born to dams exposed to heat stress conditions during the first 45 days of gestation (n = 10).

### Effects of heat stress on growing goats

The results from behavior parameters at 8 months of age obtained during the heat exposure trial are summarized in Table 4. No differences were observed between GTN and GHS growing goats in any of the parameters (*P* > 0.10) during the trial. Only lying-straight showed a treatment per time interaction trend (*P* = 0.099), however, no further differences were encountered between GTN and GHS animals neither the basal nor the heat-stress period.

**Table 4.** Behavioral and postural average expression (%) of growing goats over the basal period and after the heat-challenge period.

| Item | Treatment<sup>1</sup> | | SED<sup>2</sup> | Effect ($P$-value) | | |
| --- | --- | --- | --- | --- | --- | --- |
|  | GTN | GHS |  | Trt<sup>3</sup> | Week<sup>4</sup> | Trt×week |

| **Feeding behavior (%)** |  |  |  |  |  |  |
| --- | --- | --- | --- | --- | --- | --- |
| Feeding | 24.0 | 25.2 | 2.14 | 0.702 | 0.001 | 0.533 |
| Rumination | 14.1 | 16.9 | 1.50 | 0.179 | 0.001 | 0.345 |
| Drinking | 2.02 | 1.58 | 0.404 | 0.042 | 0.001 | 0.857 |
| **Non-feeding behaviors (%)** |  |  |  |  |  |  |
| Exploration | 4.46 | 4.88 | 0.814 | 0.709 | 0.001 | 0.231 |
| Grooming | 3.81 | 3.77 | 0.515 | 0.957 | 0.001 | 0.613 |
| Other | 3.07 | 2.77 | 0.443 | 0.645 | 0.003 | 0.390 |
| Resting | 41.2 | 37.9 | 2.35 | 0.312 | 0.001 | 0.361 |
| **Thermally-associated behavior (%)** |  |  |  |  |  |  |
| Open-mouth panting | 0.99 | 1.18 | 0.627 | 0.786 | 0.001 | 0.989 |
| Close-mouth panting | 41.6 | 37.0 | 3.90 | 0.448 | 0.001 | 0.502 |
| **Postures (%)** |  |  |  |  |  |  |
| Standing-walking | 1.38 | 1.55 | 1.547 | 0.644 | 0.001 | 0.985 |
| Standing-immobile | 33.6 | 35.3 | 2.64 | 0.643 | 0.001 | 0.718 |
| Lying-joint | 54.5 | 50.9 | 3.82 | 0.505 | 0.001 | 0.703 |
| Lying-straight | 4.95 | 5.48 | 2.172 | 0.859 | 0.001 | 0.099 |
| Neck extended | 0.42 | 0.23 | 0.233 | 0.736 | 0.006 | 0.249 |

<sup>1</sup> GTN, kids born to dams exposed to thermal-neutral conditions during the first 45 days of gestation (n = 8); GHS, kids born to dams exposed to heat stress conditions during the first 45 days of gestation (n = 8).
<sup>2</sup> Standard error of the difference.
<sup>3</sup> Trt, treatment effect (GHS vs GTN).
<sup>4</sup> Basal period corresponded to the first week at thermal-neutral conditions and heat-stress (HS) period corresponded to the following three weeks.

All parameters were affected after the heat-stress challenge regardless of the treatment (GTN vs. GHS) as shown in Fig 1. Feeding, exploration and grooming behaviors were reduced immediately after the heat challenge (week 2) and remained low compared to the basal period (*P* < 0.001) in both, GTN and GHS goats. Rumination was also lower after the heat challenge, but it started to recover towards the end of the experiment although never reached basal thermal-neutral values (week 4; *P* < 0.001). Drinking behavior also increased dramatically during the first week of exposure to heat (*P* < 0.001), but returned to initial values at the end of the experiment. Resting also increased progressively throughout the exposure to heat stress although did not reach basal values by the end of the experiment.

**Fig 1.**
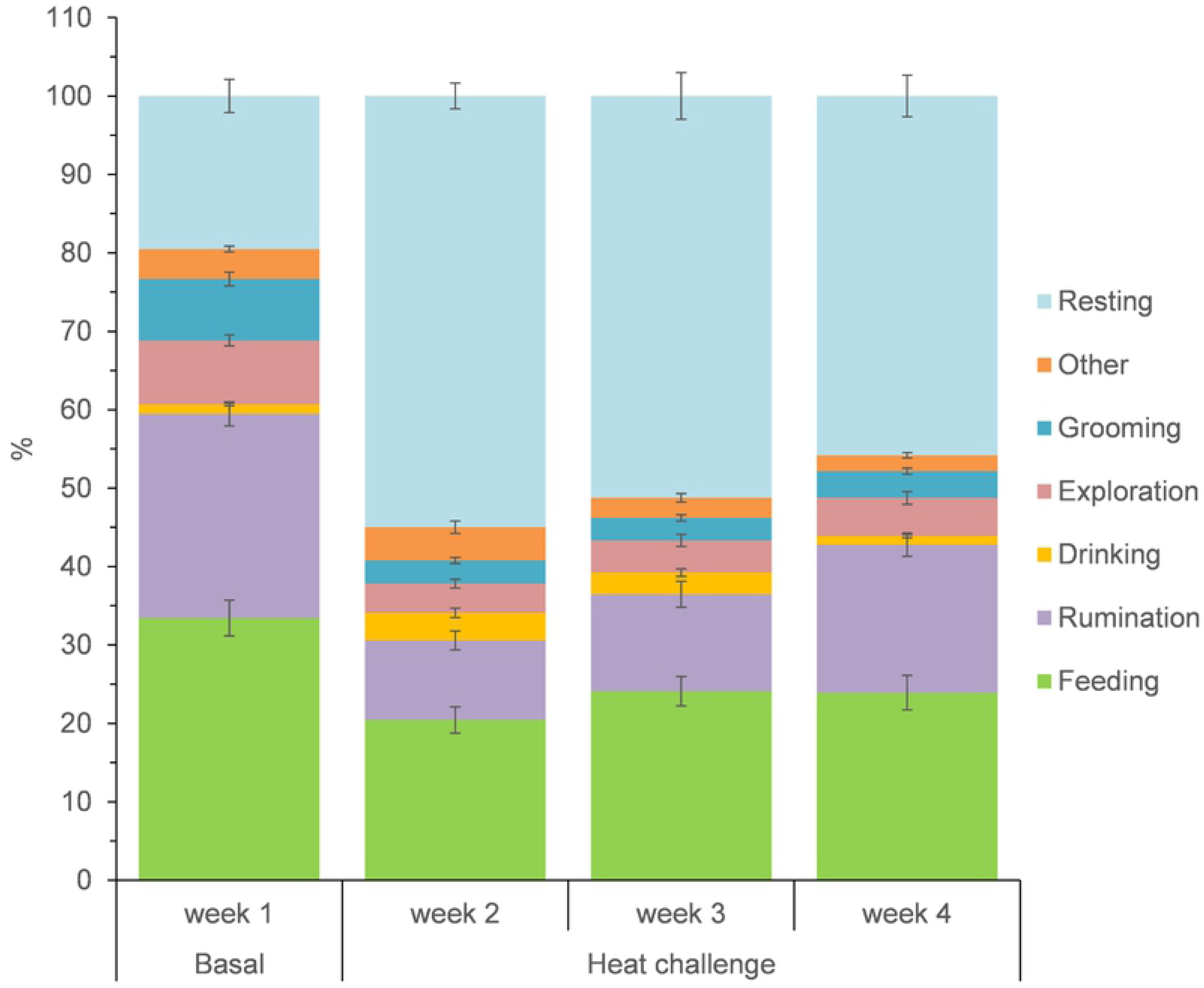
Activity behavior average expression (%) of growing goats over the basal thermal-neutral period (week 1) and during the heat-challenge period for 3 weeks (weeks 2 to 4). Bars indicate standard error.

Postural behaviors averages are presented in Fig 2. Animals were lying more frequently during the heat challenge (*P* < 0.01), predominantly with legs joint, detrimental to standing that was less observed over the heat challenge (*P* < 0.01).

**Fig 2.**
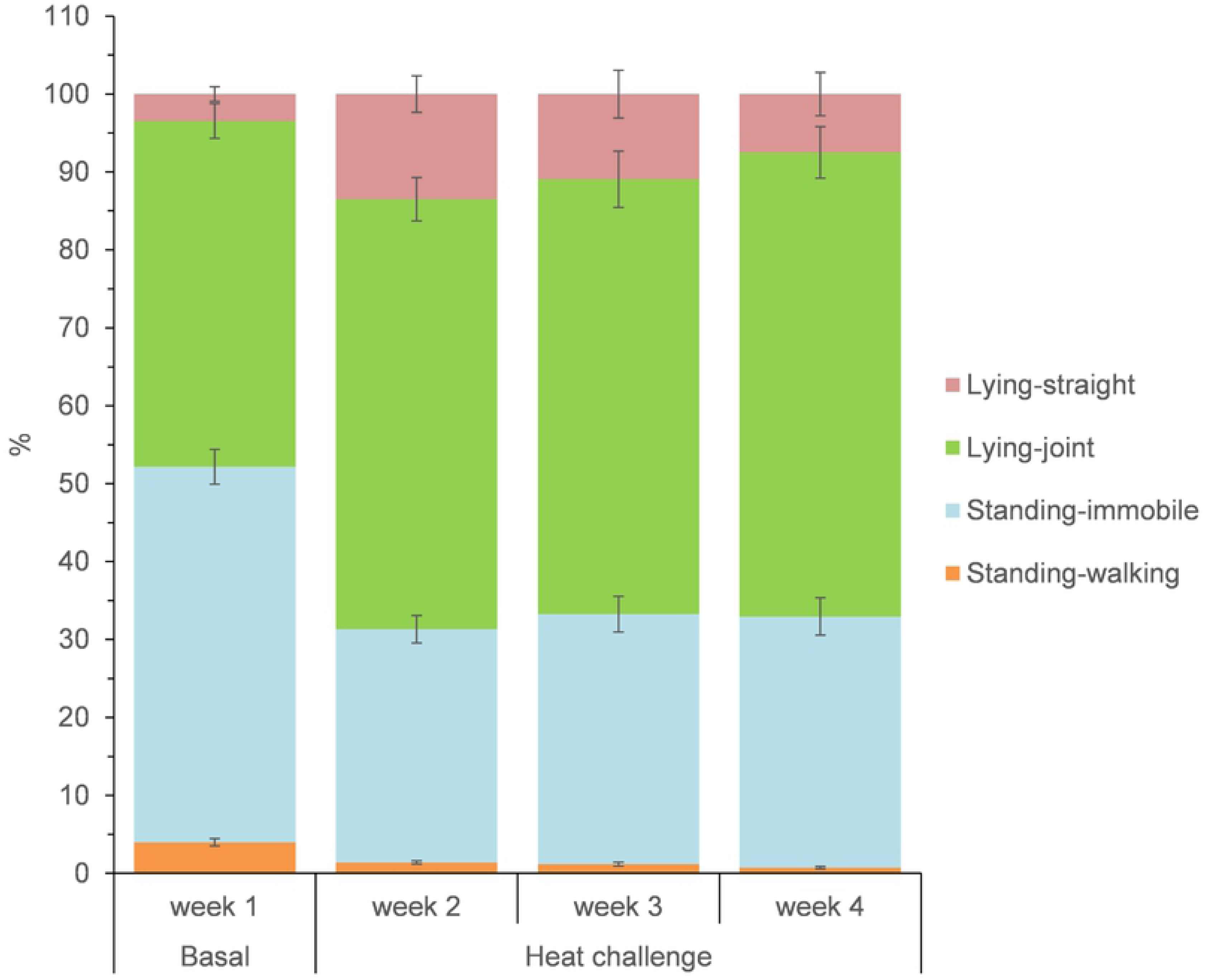
Posture average expression (%) of growing goats over the basal thermal-neutral period (week 1) and during the heat-challenge period for 3 weeks (weeks 2 to 4). Bars indicate standard error.

Additionally, as presented in Fig 3, the neck was extended more frequently compared to the basal week during the heat challenge (*P* = 0.018), as well as animals started to experience close-mouth panting after being exposed to heat and reduced this behavior progressively towards the end of the trial. Additionally, open-mouth panting was highest at week 1 of HS and disappeared at week 3.

**Fig 3.**
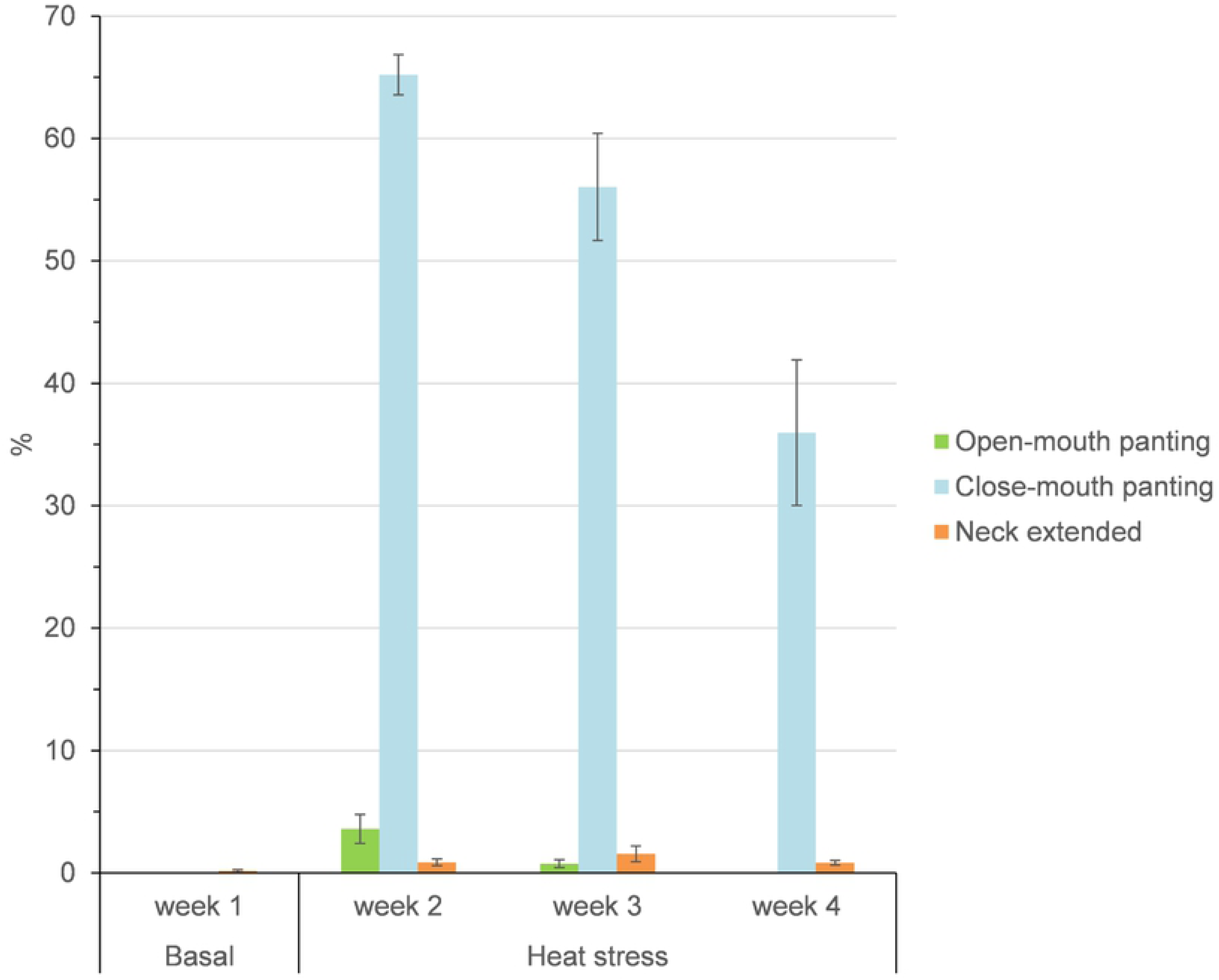
Thermally-associated behavior average expression (%) of growing goats over the basal thermal-neutral period (week 1) and during the heat-challenge period for 3 weeks (weeks 2 to 4). Bars indicate standard error.

## Discussion

In the presented study the effect of prenatal stress by exposing dairy goats to heat during mating and early pregnancy was evaluated. We aimed to investigate whether gestational exposure to heat had an impact on the development of the offspring lately in growing stages. The heat stress (HS) response was confirmed, by the evaluation of physiological parameters as well as feed intake. In this sense, GHS goats showed higher rectal temperature (+0.68°C) and respiration rate (+76 breaths/min) and a reduction by 15% in feed intake compared to GTN ones. These findings agree with previous results obtained by other authors in goats exposed to similar HS conditions [21-23]. As physiology parameters are the most reliable signs to evaluate the severity of heat stress in goats [24], goats in the present study showed clear signs of stress as a response to the heat challenge.

Although in the present study goats were mated under heat exposure, all could be effectively fertilized. Moreover, the initial hypothesis claimed that heat stress during early gestation might further affect gestation and the development of the offspring postnatally. In this regard, the most relevant outcomes were the shortening of the gestation duration of GHS goats by 3 days and the reduction of birth weight of GHS kids compared to GTN kids. Although in our study the association between the gestation length and the birth weight could not be confirmed, there is sufficient body of research that has confirmed this link in the past in sheep [25] and in cows [13,26]. These authors suggested that less time of gestation could lead to a reduction in the contributions of nutrients from the mother to the fetus. Actually, in the last 2 months of the pregnancy period is when the greatest growth of the fetus has been described in cattle (60% of the weight at birth) [27] what could partially explain the lower birth weight in GHS kids. In fact, both the shortening of the pregnancy and thus, the derived prematurity of the animals, and the thermal effect could be cofounded. Nonetheless, it is worth to mention that these associations might occur at least after the exposure to heat during the late gestation and in our case, BW differences were negligible at 35 days of life between GHS and GTN kids, which suggests that kid goats compensated the loss of fetal body growth after birth.

Nevertheless, other authors [28], were not able to demonstrate differences in the duration of pregnancy in cows exposed to heat stress at the end of pregnancy, but still the birth weight of calves born from heat stressed cows was lower in relation to its counterpart, suggesting that the reduction in the duration of pregnancy itself is not solely responsible for the reduction of birth weight, but there may be other biological changes occurring during heat stress response that affect birth weight. Heat stress during pregnancy is actually associated with poor placental development and lower blood flow, which may result in less nutrient flow to the fetus [27,29]. Additionally, Zhu et al. [16] reported that nutrient restriction of beef cows during the first third of gestation period resulted in reduced placental development and fetal weights. Hence, a reduction of nutrient supply during the first third of gestation (less feed intake by GHS) could result in impaired placental function, which negatively affects growth during gestation and contributes to lower birth weight.

In the present study we also compared the behavioral reaction to novel environments of female kids (around 1 month of life) prenatally exposed to heat stress during the mating and first 45 days of gestation. In a first approach, we implemented two tests, arena test (AT) and novel object test (NOT), in order to assess the behavioral reactivity of the kids to a new environment and an unfamiliar object, respectively. The results showed mild changes in the behavioral response of kids previously exposed to heat stress *in utero*. During the AT, GHS kids showed a reduction in the number of sniffing events in the arena. When kid goats were exposed to novel object test (NOT), a reduction in exploratory behavior (i.e. sniffing events) was also confirmed but these differences disappeared when kid goats were exposed again to NOT at 3 month of life. These results contrast with those from Roussel et al. [30], who found that kids, coming from goats under transport stress, explored (i.e. sniffing) the new environment more often than control kids. Some behavioral indicators such as immobilization, a reduction in explorative behavior and reactivity towards humans, have been related to fear [31-33]. At hormonal level, these changes have been associated to changes in the hypothalamic-pituitary-adrenal (HPA) axis [4] causing an elevation of cortisol in the maternal circulating blood during the fetal development [30,34]. Although we did not carry out measurements of cortisol in goats nor kids to confirm this casuistic, it is worth to mention that most of the development of the neural system takes place during the latter phases of gestation, and in our study, goats were not exposed to heat stress during late gestation. Thus, this could be a reason why the differences found in our animals were not as consistent as previous reports.

In the long-term scenario, the effects of prenatal heat stress on kids were followed up to growing age around five months of life, whereby the behavior was assessed by scan-sampling before and after heat exposure in order to elucidate whether prenatal heat stress had any effect on the adaptive capacity. Resting and drinking increased dramatically during the first week of heat exposure. Lying and drinking are considered as ideal biological markers for assessing the severity of the heat stress response [24]. Similarly, exploratory, grooming, and feeding behaviors declined throughout the entire period of heat exposure. Also, rumination, an essential component of the ruminant behavior that is also used as an indicator of stress and anxiety [35,36], was reduced. These activities were also accompanied by changes on the posture of animals, spending more time lying during heat exposure, mainly with legs drawing into the body, and consequently less time standing. Lying and inactivity are common expressions observed after high temperature exposition as a strategy to dissipate heat and spare energy in addition to reduce feeding [37,38]. Thus, according to our results, grown kid goats triggered a stress response when first exposed to heat. However, the fact that lying and drinking were gradually decreased afterwards suggests that animals progressively adapted to the rise of temperature.

In the same line of the results obtained in the arena tests that disappeared with time, most of the behavioral parameters assessed did not differ between GTN or GHS goats neither before nor after the heat challenge (no significant interaction between treatment and week). Only a tendency was observed for lying with straight legs (*P* = 0.099). Akbarinejad et al. [39] could not demonstrate changes in the adaptation capacity after submission to heat stress at first, second and last third of gestation of cows. In this sense, most of the works cited evaluated heat stress during late gestation, observing most of the alterations. Thence, although it seemed that the offspring could have been affected at birth, later results would suggest that HS during early gestation would not affect the offspring per se out of the gestational capability of dams. Because dams were effectively stressed according to physiological measurements, heat stress during the early period of the embryo development (1 to 45 days after mating) may not induce effects on the adaptive capacity of the offspring.

## Conclusions

Heat stress during the period of mating until the first 45 days of gestation in dairy goats reduced the duration of pregnancy and the birth weight of kids. The behavioral response of kid goats to a novel environment and objects was altered by in uterus heat stress. The exposure of the fetus to the stress response of the mother (i.e., heat stress) can modify its ability to respond to other types of stress (e.g., environmental stress) in the early postnatal life. Nonetheless, in the conditions of this study (heat duration and intensity and gestation stage) such impact disappeared towards the adult life of the animals with no differences in adaptability to heat stress.

## Supporting information

**S1 Fig. Picture of the experimental facilities**. (A) Capture of the recording for the arena test (AT). (B) Capture of the recording for the novel object test (NOT).

**S1 Table. List of behavioral and postural parameters recorded by scan-sampling during the heat-challenge experiment in the growing goats**. These parameters are drawn from the Welfare Assessment Protocol for Goats [18].

## Acknowledgements

The authors are also grateful to the team of SGCE (Servei de Granges i Camps Experimentals) of the UAB for the care of the animals.

## Author Contributions

**Conceptualization:** Ahmed A. K. Salama, Gerardo Caja, Xavier Such.

**Data curation:** Wellington Coloma, Nabil Mehaba.

**Formal analysis:** Wellington Coloma, Pol Llonch.

**Funding acquisition:** Ahmed A. K. Salama, Gerardo Caja.

**Investigation:** Wellington Coloma, Nabil Mehaba, Xavier Such, Gerardo Caja, Ahmed A. K. Salama.

**Methodology:** Wellington Coloma, Ahmed A. K. Salama, Xavier Such.

**Project administration:** Ahmed A. K. Salama.

**Supervision:** Ahmed A. K. Salama, Xavier Such.

**Visualization:** Wellington Coloma.

**Writing – original draft preparation:** Wellington Coloma.

**Writing – review & editing:** Pol Llonch, Ahmed A. K. Salama.

## Data availability statement

All relevant data are within the paper and its Supporting information files.

## Funding

This work is part of a research project funded by the Spanish Ministry of Economy and Competitivity (Programa I+D+i orientada a los retos de la sociedad; Project AGL2013-44061-R), and supported by the pre-doctoral FI grant from the Agency for Management of University and Research grants (Catalonia, Spain) awarded to NM.

## Competing interests

Authors declare no competing interests.

